# Whole Genome Analysis of 132 *Clinical Saccharomyces cerevisiae* Strains Reveals Extensive Ploidy Variation

**DOI:** 10.1101/044958

**Authors:** Yuan O. Zhu, Gavin Sherlock, Dmitri A. Petrov

## Abstract

Budding yeast has undergone several independent transitions from commercial to clinical lifestyles. The frequency of such transitions suggests that clinical yeast strains are derived from environmentally available yeast populations, including commercial sources. However, despite their important role in adaptive evolution, the prevalence of polyploidy and aneuploidy has not extensively analyzed in clinical strains. In this study, we have looked for patterns governing the transition to clinical invasion in the largest screen of clinical yeast isolates to date. In particular, we have focused on the hypothesis that ploidy changes have influenced adaptive processes. We sequenced 145 yeast strains, 132 of which are clinical isolates. We found pervasive large-scale genomic variation in both overall ploidy (34% of strains identified as 3n/4n) and individual chromosomal copy numbers (36% of strains identified as aneuploid). We also found evidence for the highly dynamic nature of yeasts genomes, with 35 strains showing partial chromosomal copy number changes and 8 strains showing multiple independent chromosomal events. Intriguingly, a lineage identified to be baker/commercial derived with a unique damaging mutation in *NDC80* was particularly prone to polyploidy, with 83% of its members being triploid or tetraploid. Polyploidy was in turn associated with a >2x increase in aneuploidy rates as compared to other lineages. This dataset provides a rich source of information of the genomics of clinical yeast strains and highlights the potential importance of large-scale genomic copy variation in yeast adaptation.

## INTRODUCTION

Yeast species classified under the *Saccharomyces sensu stricto* species complex, in particular *S. cerevisiae*, have played vital cultural and economic roles in human history (see Sicard and Legras 2011 for review). Indeed, yeast is probably one of the first species domesticated by man. Ancient evidence for the role of *Saccharomyces* in fermentation dates as far back as the Neolithic period, including wine jars dating to 7000 B.C. China (McGovern et al. 2004), 5400-5000 B.C. Iran (McGovern et al. 1997), and ~3150 B.C. Egypt (Cavalieri et al. 2003). Another Egyptian study revealed some of the oldest evidence for the active use of yeasts in bread and beer making around 1300-1500 B.C. (Samuel 1996). Yeast are widely used in industrial applications, and yeast fermentation has been harnessed in the development of biofuel technologies that produce ethanol as a renewable source of energy (e.g. Farrell et al. 2006).

With extensive human contact over nearly 7000 years, it is not surprising that *S. cerevisiae* is often found as an asymptomatic human gut commensal (Zerva et al. 1996; McCullough et al. 1998; Posteraro 1999; Salonen et al. 2000; Erdem et al. 2003). However, it has become clear that not all human-yeast interactions are harmless. *S. cerevisiae* has been identified as the causal invasive species in fungal infections with increasing frequency, often but not necessarily associated with the use of *S. cerevisiae var. boulardii* (Perapoch et al. 2000; Lherm et al. 2002; Wheeler et al. 2003; Munoz et al. 2005; Enache-Angoulvant and Hennequin 2005; de Llanos et al. 2006a, b). Thus far, *S. cerevisiae* has been estimated to be responsible for up to 3.6% of all known cases of fungemia, the most severe clinical manifestation of fungal infections (Piarroux et al. 1999; Smith et al. 2002; Munoz et al. 2005). It is also known to cause or be associated with a multitude of other symptoms that contribute to poor medical outcomes, including pneumonia, peritonitis, and liver abscesses (Dougherty and Simmons 1982; Aucott et al. 1990; Doyle et al. 1990; Mydlik et al. 1996; Munoz et al. 2005). In extreme cases, yeast appears to be the direct cause of mortality, usually by inducing sepsis (Hennequin et al. 2000; Piarroux et al. 1999). Because of such reports, *S. cerevisiae* is now considered a human pathogen, though one of relatively low virulence (Malgoire et al. 2005).

Unlike species that are fully adapted to a pathogenic lifestyle, it is unclear what factors play a role in the opportunistic virulence of a usually benign commensal. There is clear epidemiological evidence supporting impaired host immunity as an important factor. For example, a large proportion of identified cases were of low virulence and occurred in patients who were immuno-compromised, had underlying medical conditions, underwent recent antibiotic treatments, used intravenous catheters, or a combination thereof (Enache-Angoulvant and Hennequin 2005; Munoz et al. 2005). Infections appear to occur frequently through inadvertent exposure to environmental sources such as contaminated catheters, probiotic treatments, or close contact with other infected individuals (Perapoch et al. 2000; Lherm et al. 2002; Cassone et al. 2003; de Llanos et al. 2004). This is supported by evidence that *S. cerevisiae* cannot cross the epithelial barrier *in vitro* in the absence of epithelial compromise (Perez-Torrado et al. 2012).

However, even given fortuitous exposure and host condition, there is some evidence that at least two inherent phenotypic traits of yeast strains, the ability to survive at high temperatures and the propensity for pseudohyphal formation, are highly associated with virulence (McCusker et al. 1994). Virulent isolates are more likely to produce multiple distinct colony phenotypes as compared to avirulent isolates, which may contribute to their ability to survive in complex host environments (Clemons et al. 1996). A study using mice infected with clinical and non-clinical strains showed that some clinical strains appeared more virulent than non-clinical strains, but virulence phenotypes generally overlap between the two groups, suggesting that pre-existing virulence phenotypes play a complementary role to opportunistic exposure (Clemons et al. 1994). It must be noted that another study found no appreciable differences between clinical and non-clinical strains (Klingberg et al. 2008), although this result does not contradict the possibility that a subset of ‘safe’ commercial yeast strains are phenotypically more virulent (de Llanos et al. 2011), and may be more likely to cause infections given the opportunity.

Unlike phenotypic traits, the genetic factors that may influence or drive opportunistic yeast infections, if any, have been more difficult to pin down. Some gene candidates have been reported to be associated with virulence phenotypes (Muller et al. 2011; Strope et al. 2015), but most such studies were conducted on haploid derivatives from clinical isolates, whereas a microsatellite study showed that clinical yeasts are often highly aberrant in their ploidy (Muller and McCusker 2009a). This variability in ploidy and chromosomal copy number is unsurprising yet important to our understanding of yeast adaptation. Historically, yeast has had to adapt to new, stressful and drastically different environments repeatedly in the process of domestication and commercialization. Some of the best examples of how yeast strains accomplished this come from our understanding of how *S. cerevisiae* adapted to commercial fermentation (see Dequin and Casaregola 2011 for review). Commercial yeasts are extremely tolerant of large-scale copy number changes and both bread and brewing strains are often found to have >2n genomes (Albertin et al. 2009). Certain strains of sherry yeasts have stable aneuploid chromosomes and a few ale strains have 4n alleles at several loci (Legras et al. 2007; Albertin et al., 2009). Large structural rearrangements and copy number variants are rampant and may be associated with selective forces in stressful environments (Bidenne et al. 1992; Carro et al. 2003; Wei et al. 2007; James et al., 2008; Schacharer et al. 2009; Dunn et al. 2012). Aneuploidy, whole chromosomal copy number changes that typically have large fitness effects (Sunshine et al. 2015), are also common. Ploidy changes and aneuploidy are thus two large-scale genomic changes known to result in substantial fitness effects in yeast, and both have been associated with adaptation (Selmecki et al. 2009; Pavelka et al. 2010; Rancati and Pavelka 2013; Thorburn et al. 2013; Zörgö et al. 2013; Selmecki et al. 2015). However, a large scale, whole genome survey of ploidy in clinical yeast strains has not been conducted. In order to investigate the true extent of ploidy variation and aneuploidy abundance in clinical yeast strains, we sequenced 145 strains of yeast in their natural ploidy states, the majority of which are clinical yeasts isolates, to an average genomic coverage of 80x per strain. We found that 32% of strains assayed contained more than 2 copies of each chromosome (3n-5n), and 35% contained aneuploidies at one or more chromosomes. This highlights the importance of whole genome and chromosomal copy number variation in yeast population genomics and adaptation.

## RESULTS

### High Coverage Whole Genome Sequencing

145 *S. cerevisiae* strains were cultured and sequenced in their natural ploidy states. 13 of these strains were from non-clinical sources; the remaining 132 came from clinical sources. 8 strains were recorded as recently derived from strains already present in the collection, and were expected to be genetically very similar. The clinical strains were collected from at least 12 different geographical locations and isolated from at least 14 human tissue types or procedures. Some of the strains have been previously studied in one or more earlier publications: YJM798 was one of the best-studied clinical yeast strains with a fully sequenced genome (Wei et al. 2007), 14 strains were part of a clinical yeast pathogenesis study (Clemons et al. 1994), 4 strains were part of a clinical yeast phenotypic study (de Llanos et al. 2006a), 85 strains were part of a genetic diversity by microsatellite study (Muller and McCusker 2009a), 13 strains were part of a GWAS study (Muller et al. 2011), and 45 strains were sporulated and one segregant fully sequenced in (Strope et al. 2015) [Table S1]. Unlike earlier studies, such as Liti et al. 2009 and Strope et al. 2015, we sequenced the clinical strains as is, without sporulation. While extensive heterozygosity, coupled with polyploidy and aneuploidy precludes *de novo* genome assembly, it does of course provide information on chromosomal copy number and heterozygous sites.

Library sequencing, mapping, and analysis protocols are detailed in full under Materials and Methods. Briefly, libraries were constructed for each strain, and 101 bp. paired-end Illumina reads were generated, with an intended average genome wide read depth of ~80X. As it is also known that both hybridization and introgression are not uncommon in clinical budding yeasts (Muller and McCusker 2009b; Strope et al. 2015), the fastq reads were mapped to a combined *Saccharomyces sensu stricto* genome that included sequences from *Saccharomyces cerevisiae (S.cer), S. paradoxus (S.par), S. mikatae (S.mik), S. kudriavzevii (S.kud)*, and *S. uvarum (S.uva)*. The S288C reference sequence was downloaded from the *Saccharomyces* Genome Database; all other species were downloaded from *Saccharomyces sensu stricto* Resources (www.saccharomycessensustricto.org). One strain was subsequently removed from analysis due to a likely contamination during library construction, with the vast majority of reads failing to map entirely [Methods]. The coverage depth for the other 144 individual strains ranged from 57X to 215X within the *S.cer* sequences. Reads were then deduplicated, realigned, and recalibrated following best recommended practice on the Genome Analysis Toolkit (GATK) website. Single nucleotide polymorphic sites (SNPs) were filtered by coverage depth, mapping quality, and minimum number of reads supporting variant allele. SNPs falling within known repeat sequences and telomeric regions were subsequently excluded from analysis.

### No Clinical Specific Point Mutations Found

Within *S. cerevisiae* genomic sequences, a total of 477,512 single nucleotide polymorphic sites (SNPs) were called across all strains, with individual strains carrying between 36,786 – 97,758 variants (0.3%-0.8%) as compared to S288C. Although the number of non-clinical strains was small, allele frequencies at diallelic positions could be compared between clinical and non-clinical strains. Correlation between the populations was high, with an r^2^ value of 0.9346 [Figure S1]. F_st_ values were calculated for every position, but values did not exceed 0.15, signifying that no SNPs were differentiated between clinical and non-clinical strains [Figure S2]. Thus, no further analysis to identify clinical specific alleles was performed.

### Population Phylogeny Recapitulate Earlier Studies

A neighbor-joining (NJ) tree was constructed with the newly sequenced strains as well as 35 sequences previously published in Liti et al. 2009 (Figure 1). The two NA soil strains, the three Malaysian strains, the three sake strains, the two baker strains, and the lab strains W303 and S288C cluster into independent clades, as would be expected (Strope et al. 2015). The clade consisting of YJM789, a strain isolated from lung, also included other strains isolated from lung, paracentesis fluid, and blood. Six clinical strains appeared to be closely related to Fleischmann’s baking yeast. Twenty-three clinical strains fell with four commercial yeast strains from Italy on the NJ tree (although it was unclear what precisely the commercial niche was). Four of these 23 strains were documented as clinical isolates from nearby Greece. Three clinical strains further down the tree appeared to be nearly identical to a pro-biotic strain labeled as *S. boulardii*. Ten strains were closest to two known vaginal isolates, although information regarding their sites of isolation was not available. A handful of strains fell with RM11_1A, a derivative of a natural vineyard isolate from California. Finally, a couple of clinical strains appeared most similar to baker yeast strains.

**Figure 1.**
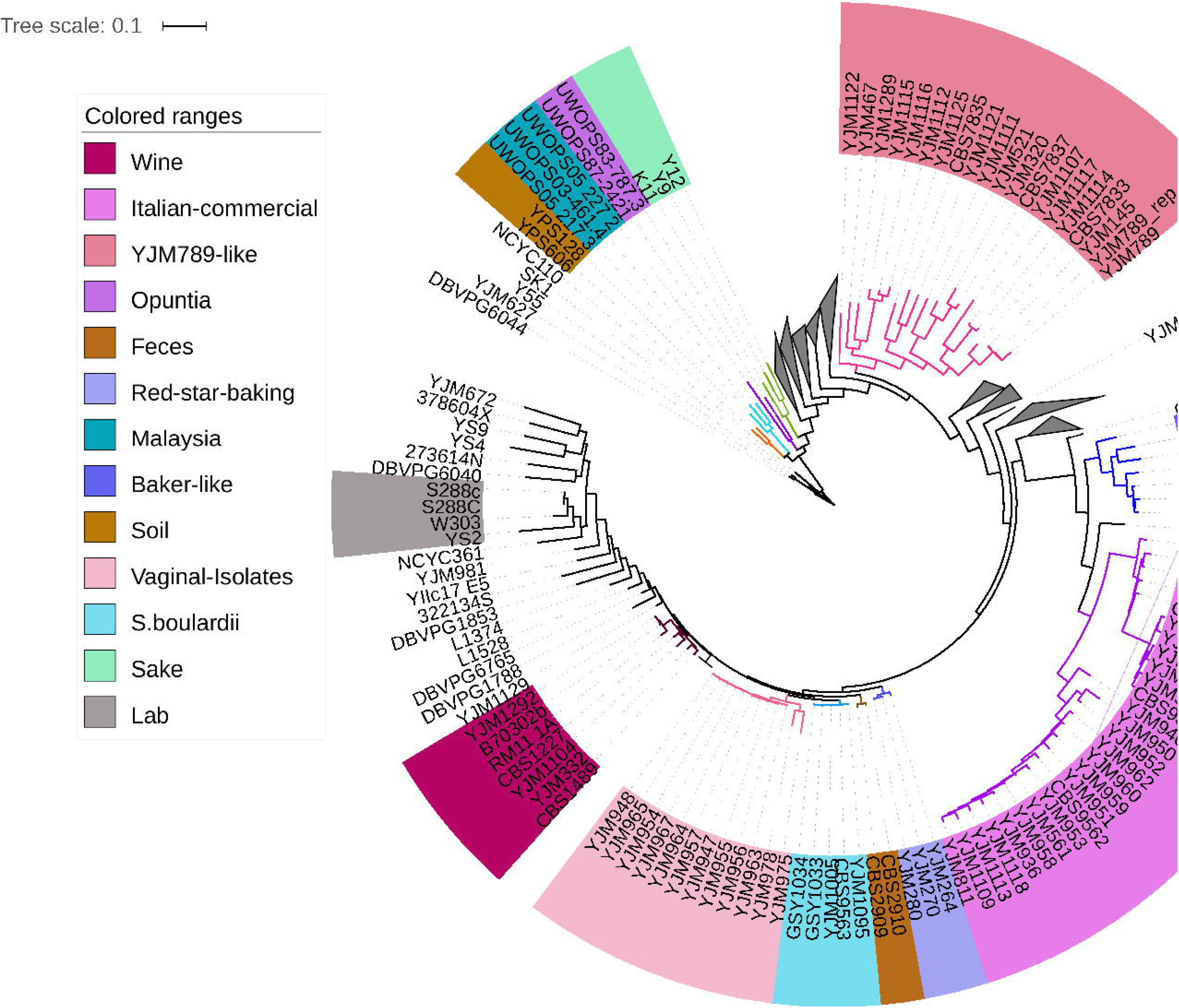
Neighbour-joining tree of 145 sequenced strains and 35 Liti et al. 2009 strains, based on SNP differences within the *S. cerevisiae* genome. Groups of strains that fall within a cluster containing non-commercial strains from a single known source are highlighted in color blocks. Clades containing only clinical strains without a known source are collapsed, with the full tree available in Figure 4.

### Population Structure Reports Multiple Origins Of Clinical Strains

Overall, the clinical strains were scattered throughout the clades of non-clinical strains on the phylogenetic tree, suggesting multiple independent origins. To obtain a more explicit grouping of ancestral source population, we used Structure (Pritchard et al. 2000; Falush et al. 2003, 2007; Hubisz et al. 2009) on the same set of SNPs used to create the NJ tree [Methods]. The clinical strains were distinguished into 5 main populations in our analysis. The non-clinical strains that corresponded to these populations included baking strains, sake/soil strains, wine strains, and unspecified commercial strains from Italy. A fifth population consisted of only clinical strains, including YJM789, similar to previous studies that found an unknown source population for clinical strains (Liti et al. 2009; Strope et al. 2015). The remainder of the clinical strains showed varying amounts of admixture from various population sources (Figure 2).

**Figure 2.**
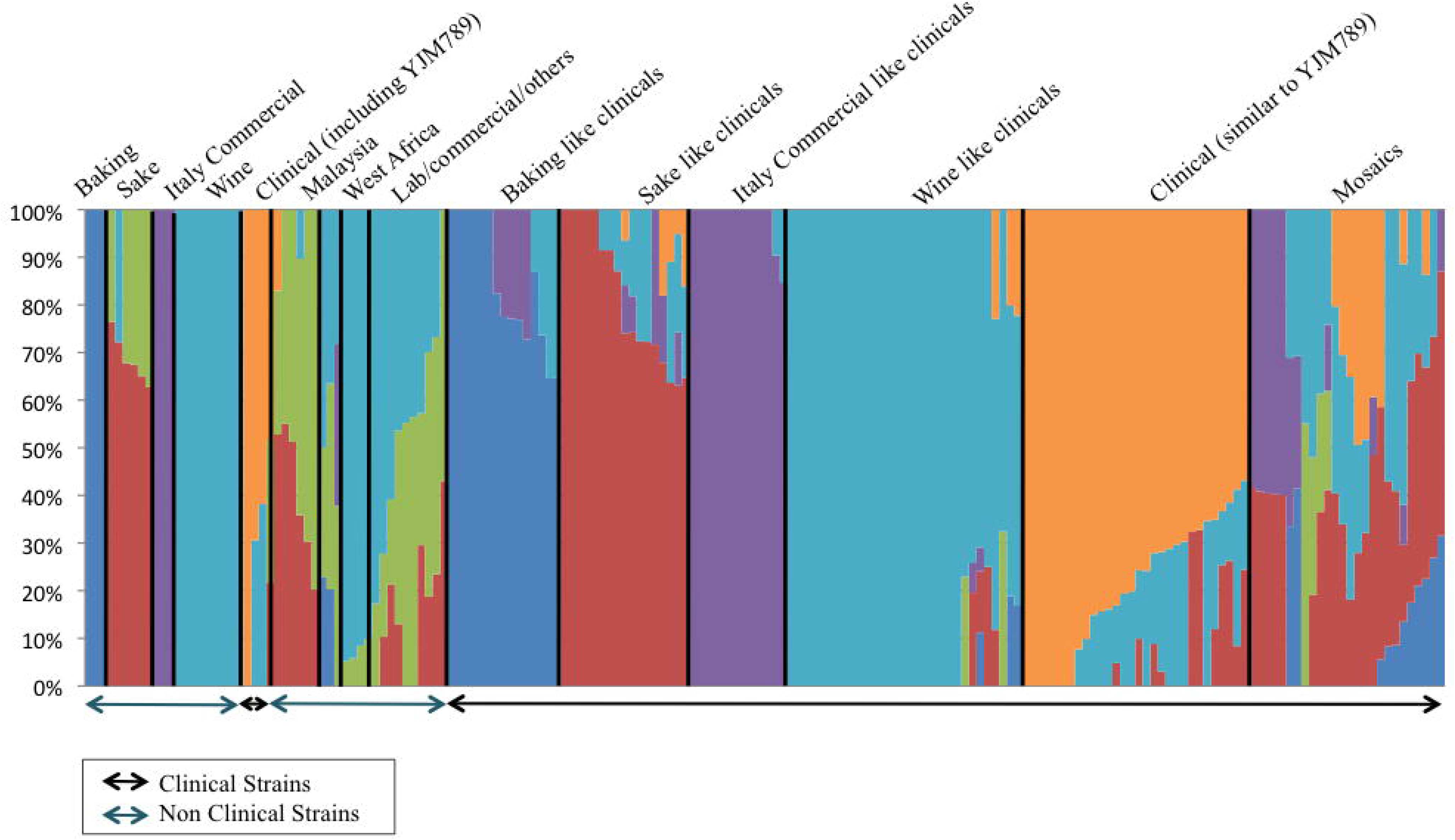
Structure assignment of source populations for all 180 strains. Structure (version 2.3.2) analysis was run with correlated frequencies, admixture model based on base pair distance from preceding SNP, 15,000 burn in period and 5,000 iterations of sampling.

### Introgressions From *S. paradoxus* and *S. kudriavzevii*

In addition to reads mapping to the *S.cer* genome, three strains also had some sequence reads that mapped to sequences outside of the *S.cer* region of the pan-genome.

YJM264 is a known hybrid (Muller 2009). Reads covered complete chromosomes from both *S.cer* and *S.kud* species. Coverage plots across the genomes showed the presence of at least one copy of all *S.kud* chromosomes except V, XI, XIV and XVI, and likely two copies of *S.kud* chromosomes IX and XV. *S.kud* chromosomes XII and XIII were partially present, with reads mapping to the first 435Kb of XII and the first 364Kb of XIII. Coverage across the *S.cer* chromosomes showed complementary patterns, including lower copy numbers for the partial segments on chromosomes XII (first 450Kb) and XIII (first 390Kb), such that combined across both genomes, there are three copies of all chromosomes. The only exceptions were chromosome V, which appeared to have 4 copies, and chromosome II, which appeared to have a segmental duplication spanning the first 200Kb [Figure S3].

CBS2909 and CBS2910 are two strains isolated from human feces that fell immediately next to each other on the phylogeny. Both strains showed highly concordant, though not perfectly identical, regions of high coverage along the *S.par* genome, ranging between 2kb-40kb in size. 818 such 1kb windows were observed in CBS2909 and 778 such 1kb windows were observed in CBS2910. This mapping pattern was not observed in any other strain. Mapping results suggest them to be mainly *S.cer* genomes with *S.par* or *S.par*-like fragments interspaced across the genome [Figure S4]. Removing regions that may be prone to mismapping artifacts (artificial coverage in >50% of all strains), 47 remaining regions consisting of 272 *S.par* genes were suspected of introgression into CBS2909 (279 genes) and CBS2910 (274 genes) [Table S2]. 261 genes were shared by both strains. Other yeast strains have also been previously found to contain introgressions from *S.par* (Strope et al. 2015). *S.par* is often found in the same habits as *S.cer*, and such results may indicate ancient hybridization (Sampaio and Gonçalves 2008; Sniegowski et al. 2002).

No other clinical strains showed significant coverage across non*-S.cer* sequences.

### Ploidy Variation Is Common In Clinical Strains

In *S. cerevisiae* strains, ploidy variation is common. For 96 of the strains that have been included in previously published work, sporulation patterns indicated that 11 strains were haploids, 56 were diploids, 15 were triploids, and 14 were tetraploids (Muller and McCusker 2009a). However, because a number of generations had elapsed since the strains were first made or studied, we assigned ploidy state to each of our sequenced strains *de novo*, using observed SNP frequency distributions within the sequence reads. Haploid strains should contain only SNP frequencies of 1; diploids should contain frequencies of both 0.5 and 1, as some SNPs will be heterozygous, while others will be homozygous, and so on (Figure 3). We note however, that homozygosing events will eliminate heterozygous SNPs, while not changing ploidy. Thus, we were unable to unequivocally rule out euploidy in homozygous strains in the absence of aneuploidy or previous published work. All 11 haploid strains matched expectations. Three of the diploid strains, YJM1104/YJM1105/YJM436, appear to have been recently homozygozed, possibly during isolation and propagation of the strain – their SNP distributions had no visible clusters of SNPs at p=0.5 as compared to other diploid strains. YJM525 and YJM954 were listed as diploid strains with profiles similar to that expected of triploids. YJM464, YJM1094, YJM1099, and YJM1114 were listed triploid strains that had SNP frequency distributions that indicated diploidy. YJM1146 was also listed as triploid but looked tetraploid. Finally, YJM950 and YJM952 were listed as tetraploids, but resembled triploids. In all 145 strains, there were 16 (11%) haploids, 83 (57%) diploids, 23 (16%) triploids, and 23 (16%) tetraploids. Most strains that were 3n or 4n fell within two clades on the phylogeny, the Italian commercial clade and the baker strain clade, suggesting a common recent ancestry (Figure 4).

**Figure 3.**
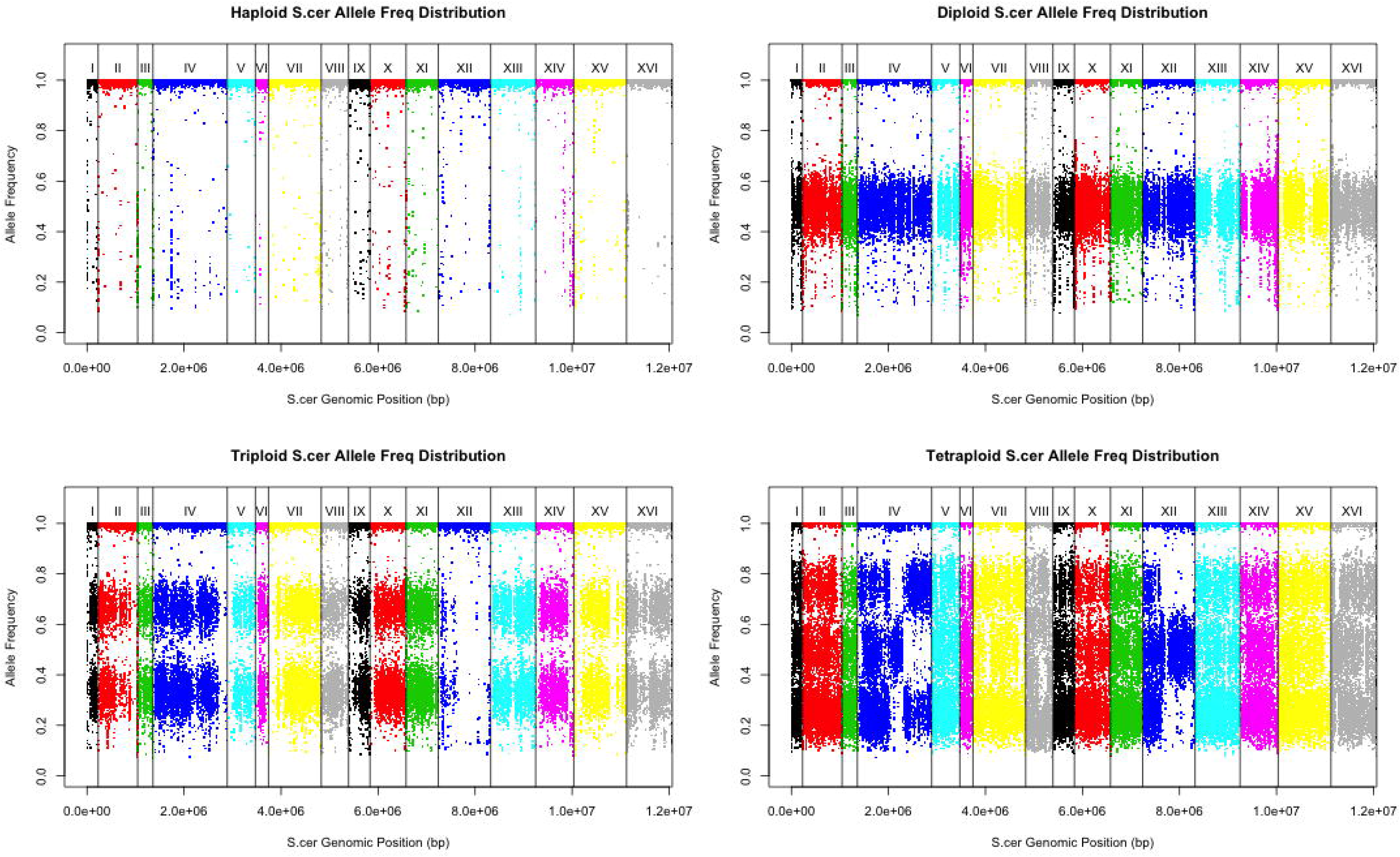
Top left) Allele frequency plot of a typical haploid with SNPs at allele frequencies at 1 Top right) Allele frequency plot of a typical diploid with SNPs at allele frequencies at 0.5 Bottom left) Allele frequency plot of a typical triploid with SNPs at allele frequencies at 0.33, 0.67, and 1 Bottom right). Allele frequency plot of a typical tetraploid with SNPs at allele frequencies at 0.25, 0.5, 0.75, and 1. Chromosomes are lined in increasing order (X-axis), with raw SNP allele frequencies along the genome plotted (Y-axis).

**Figure 4.**
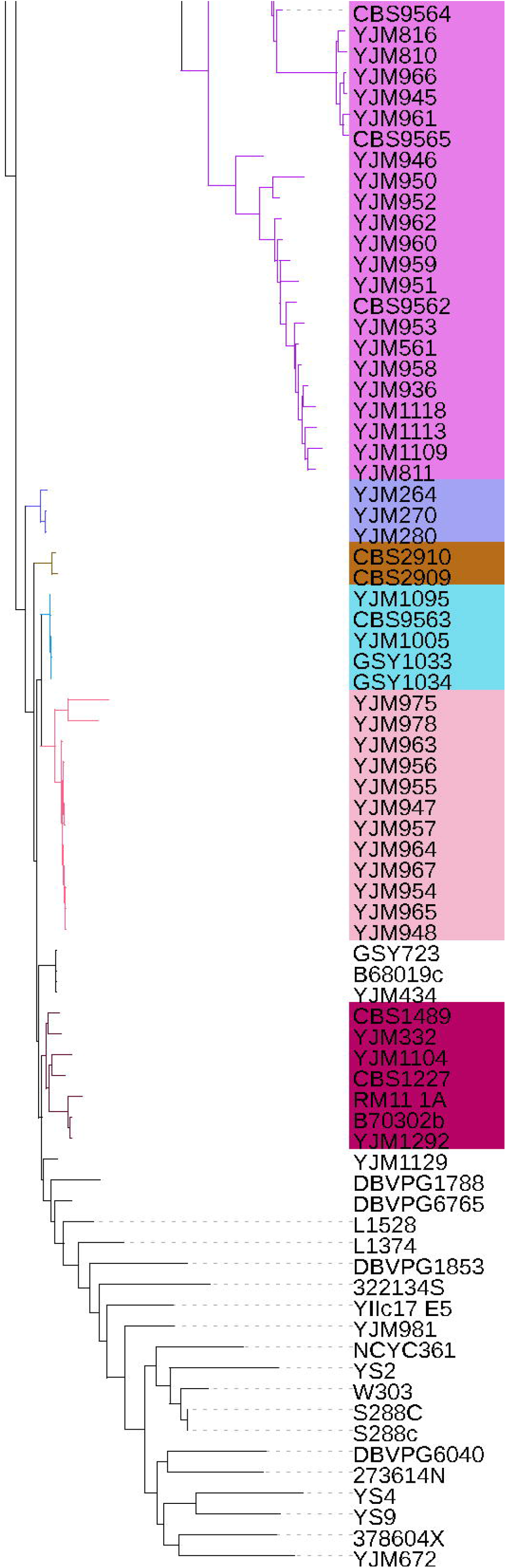
Heatmap of chromosomal copy number (0n-5n) for each of 145 strains aligned next to each strain on the phylogenetic tree. The 35 Liti 2009 strains were homozygosed and do not have chromosomal copy number information panels. The 16 chromosomes of *S.cer* are sorted in increasing order from left to right. Panel color from lightest to darkest green represent chromosomal copy number in increasing order.

### Aneuploidy Is Found In More Than One Third Of Clinical Strains Surveyed

Inconsistencies in allele frequency distributions across chromosomes within a single strain were also observed. This could have been due to recent homozygozing of individual chromosomes, or more likely due to individual chromosomes being aneuploid. We assigned the presence of aneuploidies in each strain by examining coverage for 1kb non-overlapping windows across all 16 *S.cer* chromosomes. 53 of 145 strains (36%) were observed to contain aneuploidies, a number nearly 5 times higher than the 7.5% previously observed in newly isolated segregants (Strope et al. 2015). Some chromosomes were more often observed in an aneuploid state than others, possibly correlated with chromosomal size (Figure 5). The most commonly observed whole chromosomal aneuploidy was chromosome IX, with 22 observed events across all strains, as compared to the 1-13 events observed in all other chromosomes. When the rate of observed aneuploidies was compared between strains with ploidy states that are thought to be more stable (haploids and diploids) and strains with ploidy states that are thought to be unstable (>= 3n), there was a significantly higher aneuploidy rate in the latter (1n/2n – 27.3%, >=3n – 56.5%, Fisher’s exact test *p* = 0.0009). There were also 35 instances of inconsistencies within a chromosome due to a partial chromosomal copy number change (Figure 6). Partial chromosomal copy number changes were most often seen in chromosomes III and VII. There appeared to be no correlation with chromosomal size (Figure 5).

**Figure 5.**
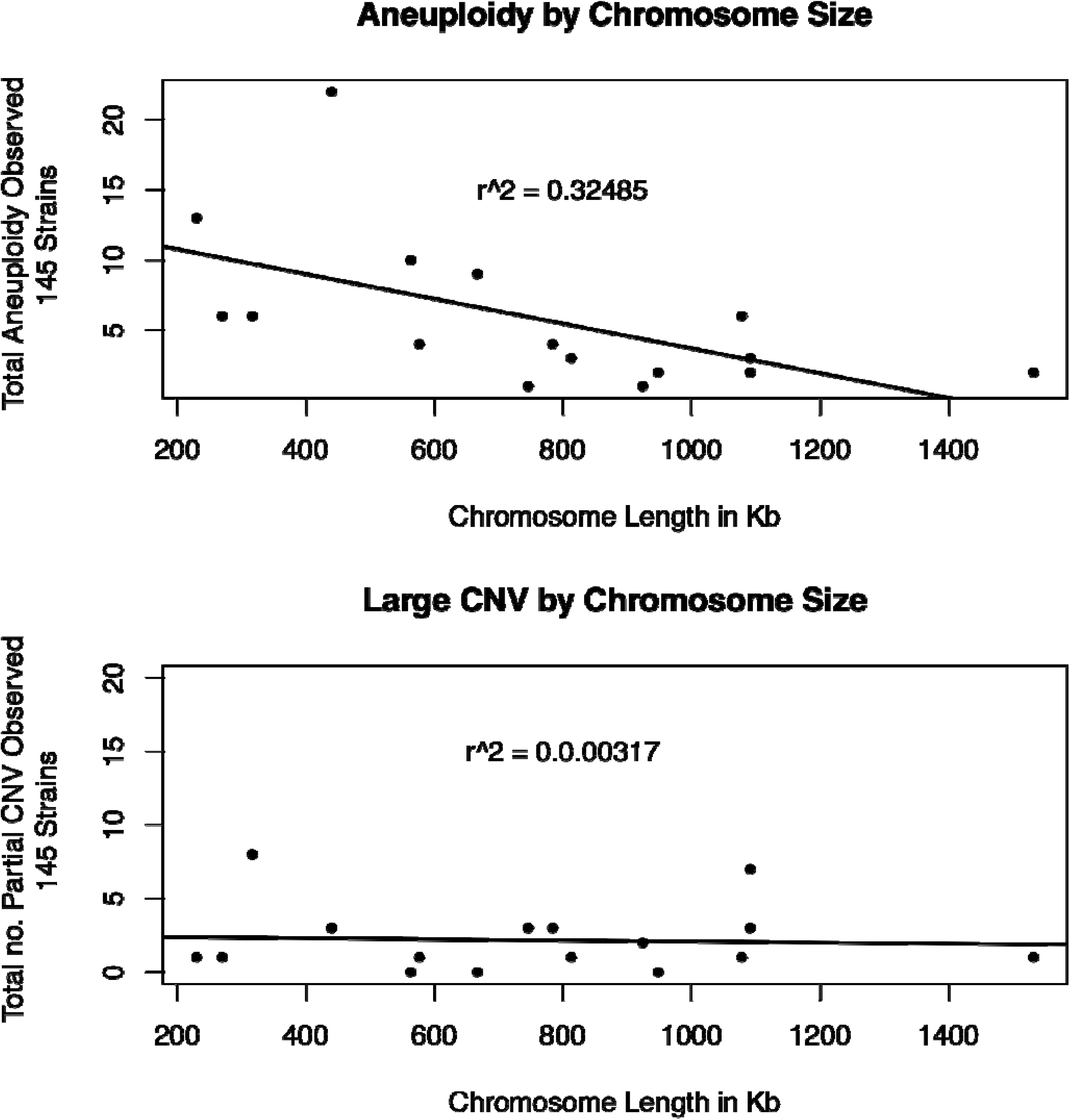
Top). Correlation between whole chromosome aneuploid events observed (Y-axis) and chromosomal size (X-axis). Bottom). Correlation between partial chromosomal copy number changes observed (Y-axis) and chromosomal size (X-axis).

**Figure 6.**
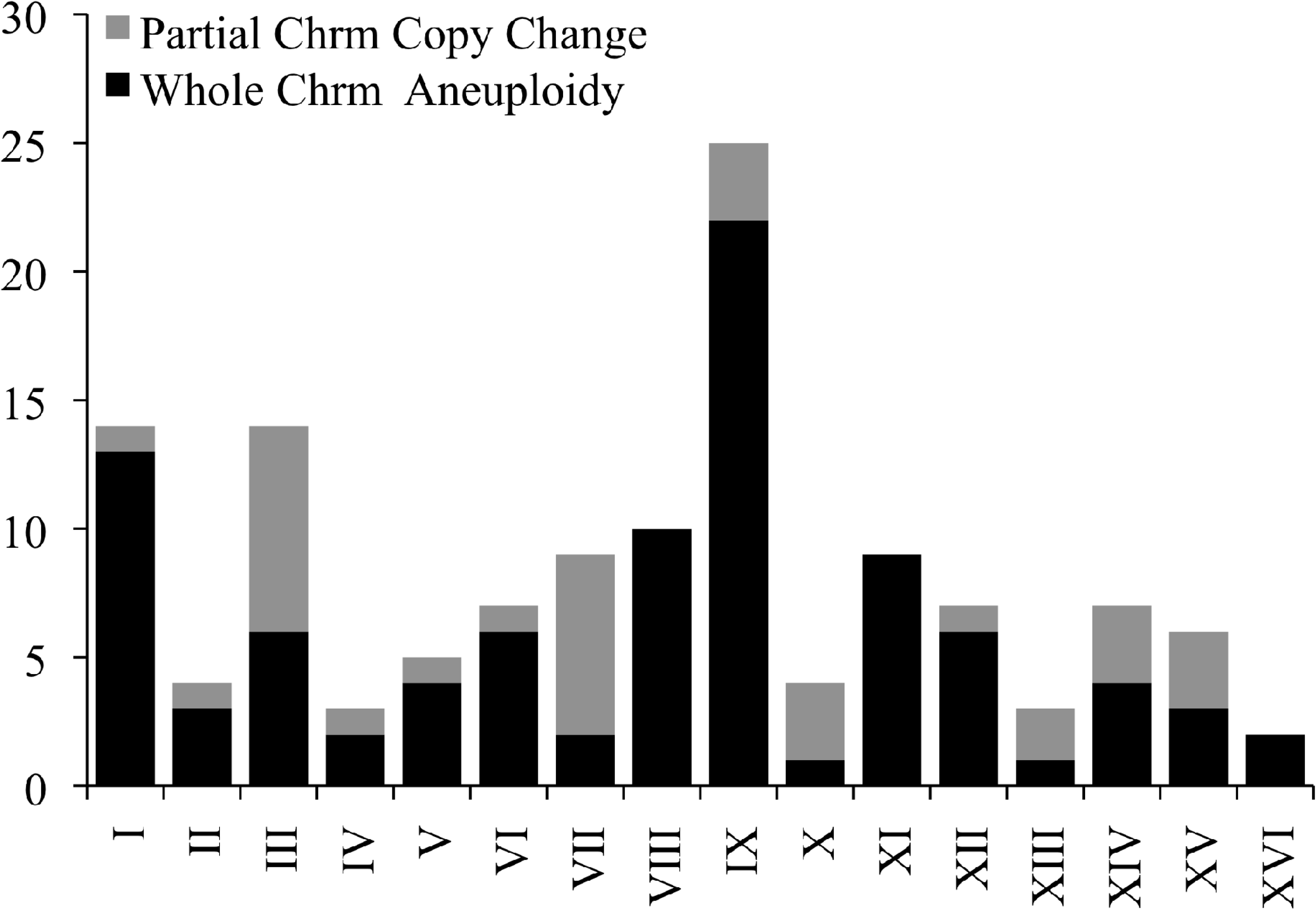
Histogram of the total number of complete and partial aneuploid events observed within all 145 sequenced strains, by chromosome.

Combined information from SNP frequency distributions and aneuploidy calls suggested that multiple chromosomal scale copy number events may be present in some strains. One diploid strain YJM1098 appeared to have a 3n aneuploid chromosome XII, but SNP frequencies also suggested that chromosome VIII has been recently homozygozed (Figure 7). Another triploid strain, YJM466, had four chromosomal copies of both chromosome VI and IX, but chromosome VI showed SNP a frequency distribution of a tetraploid, while chromosome IX showed that of a diploid (Figure 8). YJM525, YJM674, YJM810, YJM811, YJM813, and YJM815 also showed such indications of multiple events.

**Figure 7.**
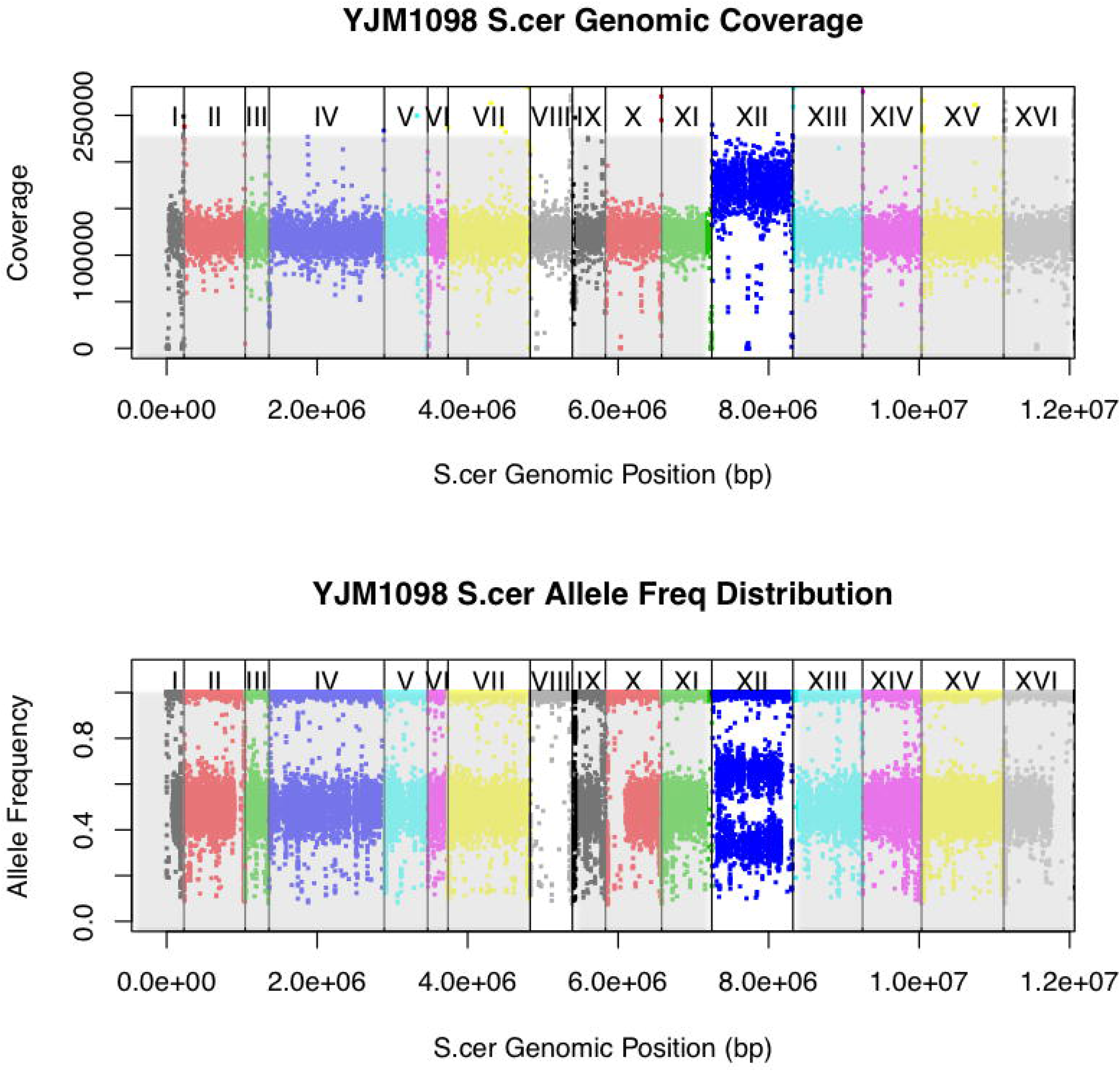
Coverage and allele frequency plots for YJM1098. Chromosomes are lined in increasing order (X-axis), with total coverage in 1kb windows [top panel] or raw SNP allele frequencies [bottom panel] plotted (Y-axis). YJM1098 showed a copy number change in chromosome XII with corresponding shift in SNP allele frequency distribution, as well as no copy number change in chromosome VIII, but a allele frequency distribution suggesting recent chromosomal homozygosing event.

**Figure 8.**
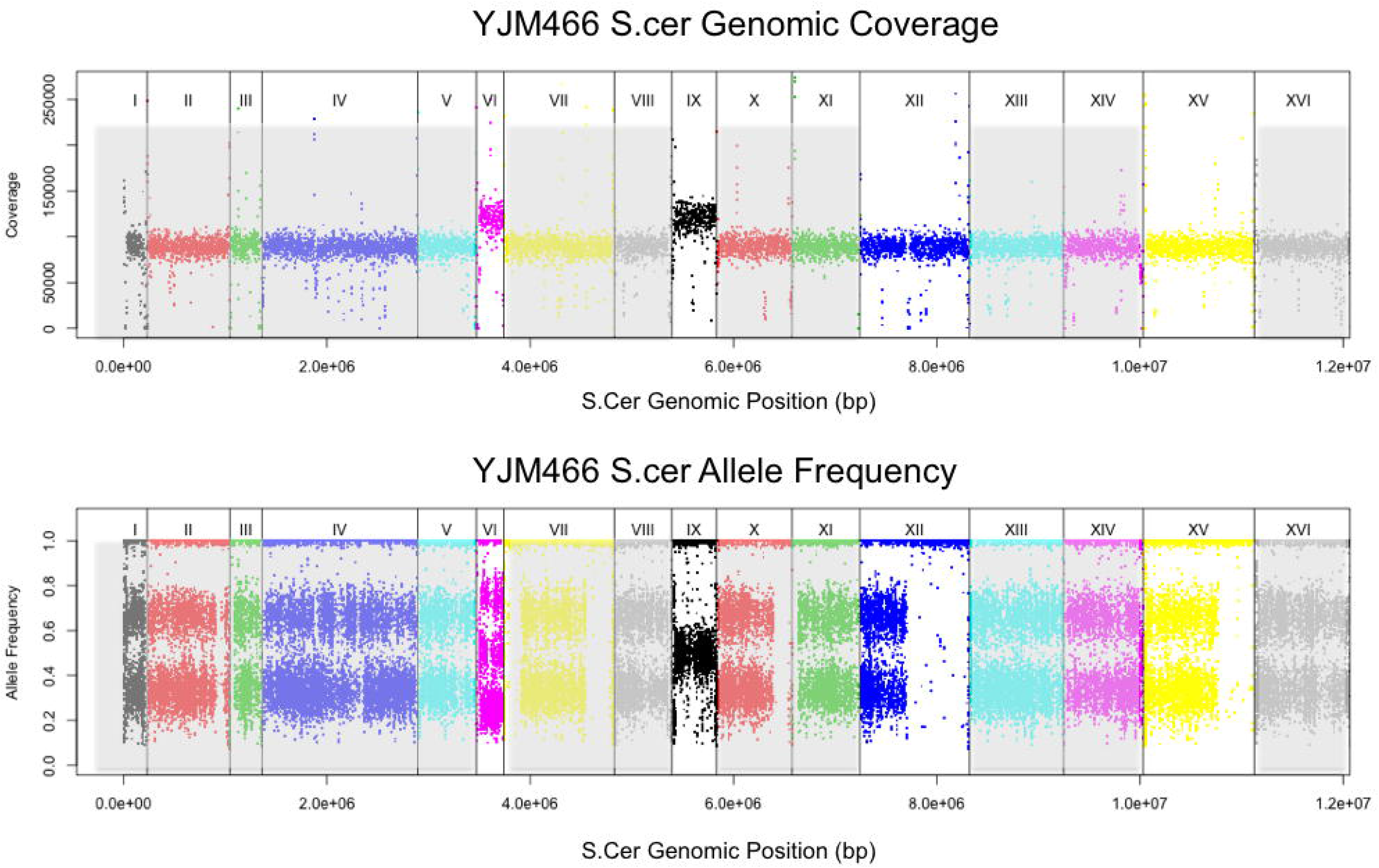
Coverage and allele frequency plots for YJM466. Chromosomes are lined in increasing order (X-axis), with total coverage in 1kb windows [top panel] or raw SNP allele frequencies [bottom panel] plotted (Y-axis). YJM466 is a triploid strain that showed a copy number increase at chromosomes VI and IX. Chromosome IX showed a corresponding shift in SNP allele frequency distribution to that of 4n, but chromosome VI showed a shift in SNP allele frequency distribution that resembles 2n.

### Damaged Copies of *NDC80*: Possible Genetic Basis of High Ploidy/Aneuploidy Rates in Baking Lineage

Strains that were polyploid or carried aneuploid chromosomes were not randomly distributed across the population. One particular clade, containing 36 members from CBS1464 to YJM811 on the phylogeny tree, contained an unusually high number of polyploid and aneuploid strains (Figure 4). 29/36 (81%) strains from the clade made up of Baker’s-like and Italian Commercial strains were 3n/4n, whereas only 17/98 (17%) of all other non-segregant strains were (Fisher’s exact test *p*=2.05e^-11^). Similarly, 24/36 (67%) of strains from the baker’s/Italian clade contained aneuploid chromosomes, whereas 32/109 (29%) of all other strains were (Fisher’s exact test *p*=1.3e^-04^).

We thus investigated if there were any mutations that might explain the high levels of aneuploidy (and ploidy) observed in this particular lineage. Hundreds of genes have been associated with genomic stability and faithful replication and segregation during the cell cycle (Ouspenski et al. 1999; Stirling et al. 2011; Cheng et al. 2012). In particular, a number of genes involved in sister chromatid separation and segregation in yeast have been well studied and characterized. We focused on the genes encoding the kinetochore proteins *NDC10* (*CBF2*), *BIR1, NDC80* (YIL144W), *CEP3*, and the chromosome segragation genes *CSE4, IPL1, SMT3* (Goh and Kilmartin 1993; Biggins et al. 2001; Cho and Harrison 2012). We used SIFT to look for damaging SNP mutations that are likely to affect the function of these genes (Methods).

Most of these genes are essential and had very few damaging mutations in the strains sampled (*BIR1*: 0 strains, *CEP3*: 3 strains, *SMT3*: 1 strain, *CSE4*: 4 strains). However, a much larger number of strains carried copies of damaging mutations in *NDC80* (41 strains), *IPL1* (57 strains), and *NDC10* (27 strains). Of these, one particular proline to glutamine mutation in *NDC80* at ChrIX:78597, mostly in the heterozygous form, was only found in the baker/commercial lineage, affected 32/36 strains, and may be part of the explanation for the high rate of aberrant chromosomal copies observed in this clade. Additionally, all 36 baker/commercial lineage strains contained damaging mutations in either *NDC80* (89%) or *IPL1* (89%), with half also carrying affected copies of *NDC10* (54%). These numbers are significantly different from strains outside the lineage, where only 29/98 strains carried any damaging mutations in these genes, and the relative frequencies are just 7% in *NDC80* (Fisher’s exact test *p*=2.2e^-16^), 26% in *IPL1* (*p*=4.5e^-11^), and 10% in *NDC10* (*p*=0.035). Strikingly, almost all other damaging mutations in *NDC80* found outside of the baker/commercial lineage were in polyploid strains, with just two exceptions in two closely related segregant strains that shared a unique mutation not found in any other strain.

### Gene Loss and Copy Number Changes In Clinical Strains

To look for gene copy number changes that may be adaptive in all clinical strains, we investigated recent gene losses and gains for verified genes in the 145 sequenced clinical strains. Genes lost or gained in a few strains may be recent sporadic events, but a common gene loss or gain found in most clinical strains could indicate an adaptive advantage.

Conservatively, genes were classified as lost only if no reads mapped to any position within the coding sequence of a gene, and the Bonferroni corrected likelihood that a gene of that length is not sequenced given each strains’ average genomic coverage was <0.05. 95 genes passed the filter [Table S3]. We focused on recent gene losses – those found only in <5% (1-7 strains) of the sampled population. The 10 most significant GO terms associated with the 52 deleted genes suggested a depletion of deletion in genes involved in cellular component organization and biogenesis [Table 1]. There was enrichment in vitamin B6 related biosynthetic and metabolic pathways, largely due to a four-gene deletion (SNZ2, SN02, SNZ3, SNO3) in one strain from chromosomes VI and XIV. These genes are found in the subtelomeric region, and are known to vary in copy number (Stambuk et al., 2009). Conversely, common gene deletions (7-86 strains) showed enrichment in deletion of genes involved in transmembrane glucoside and saccharide transporter activities [Table 1].

**Table 1.**
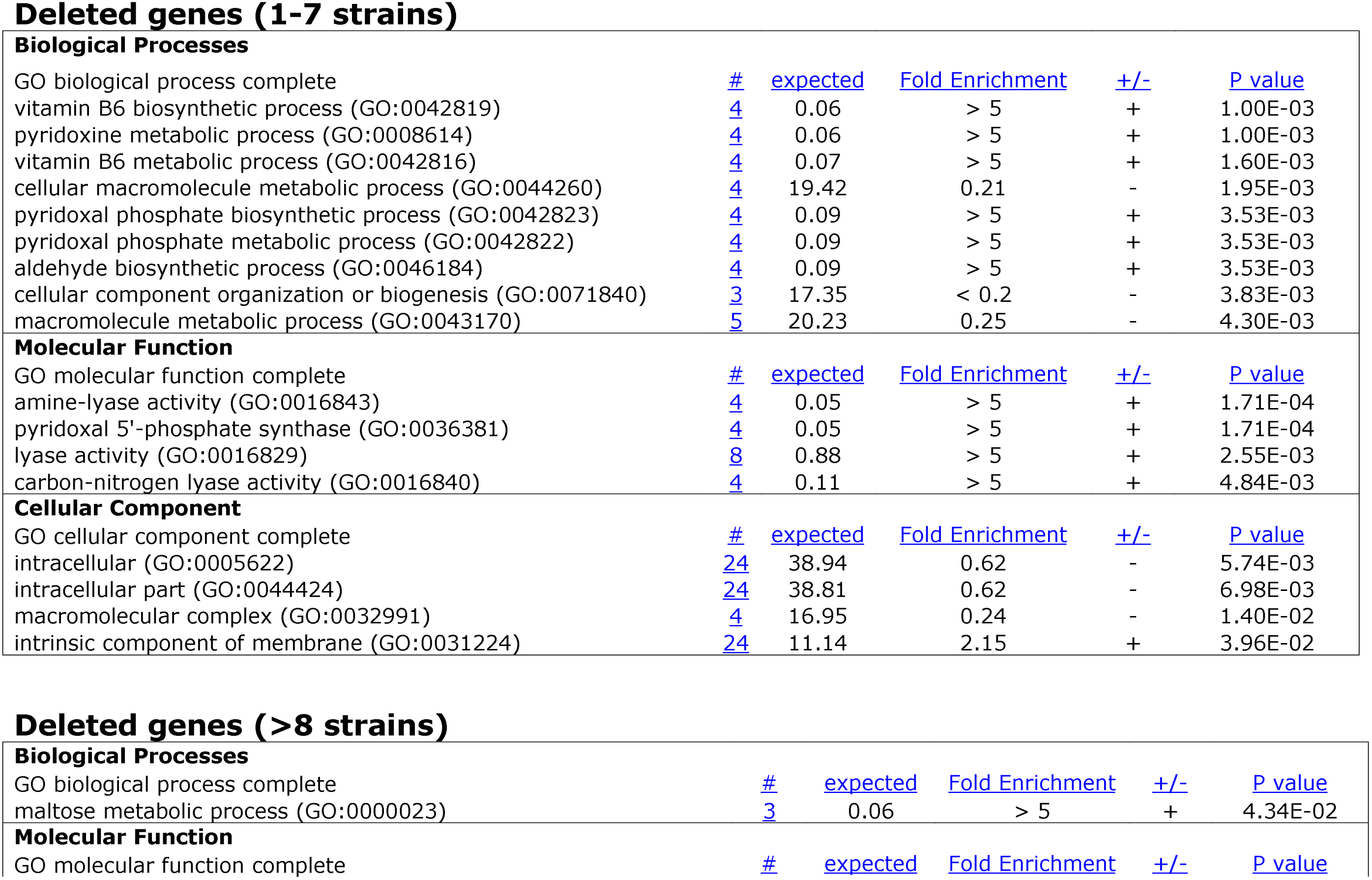
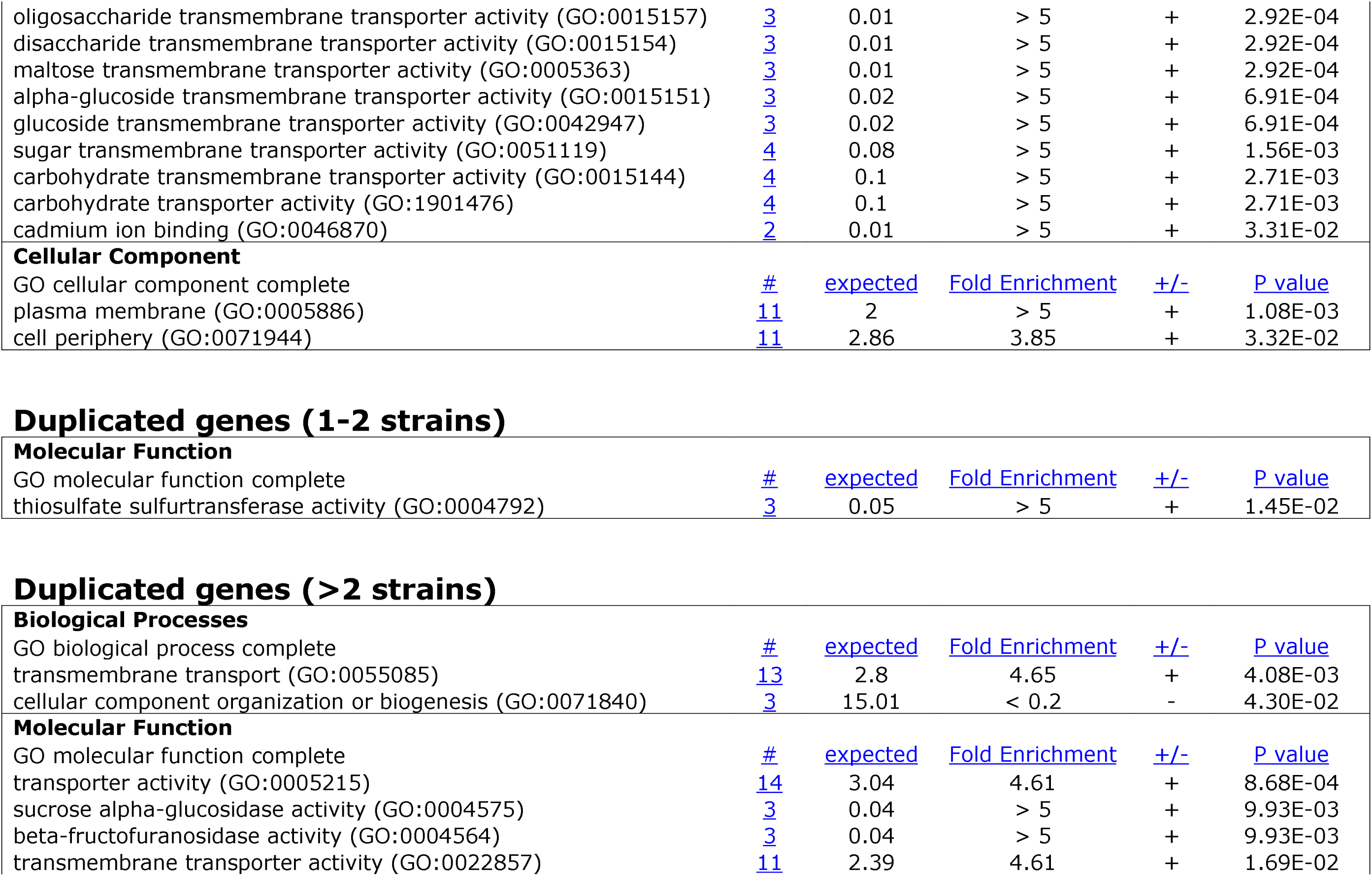
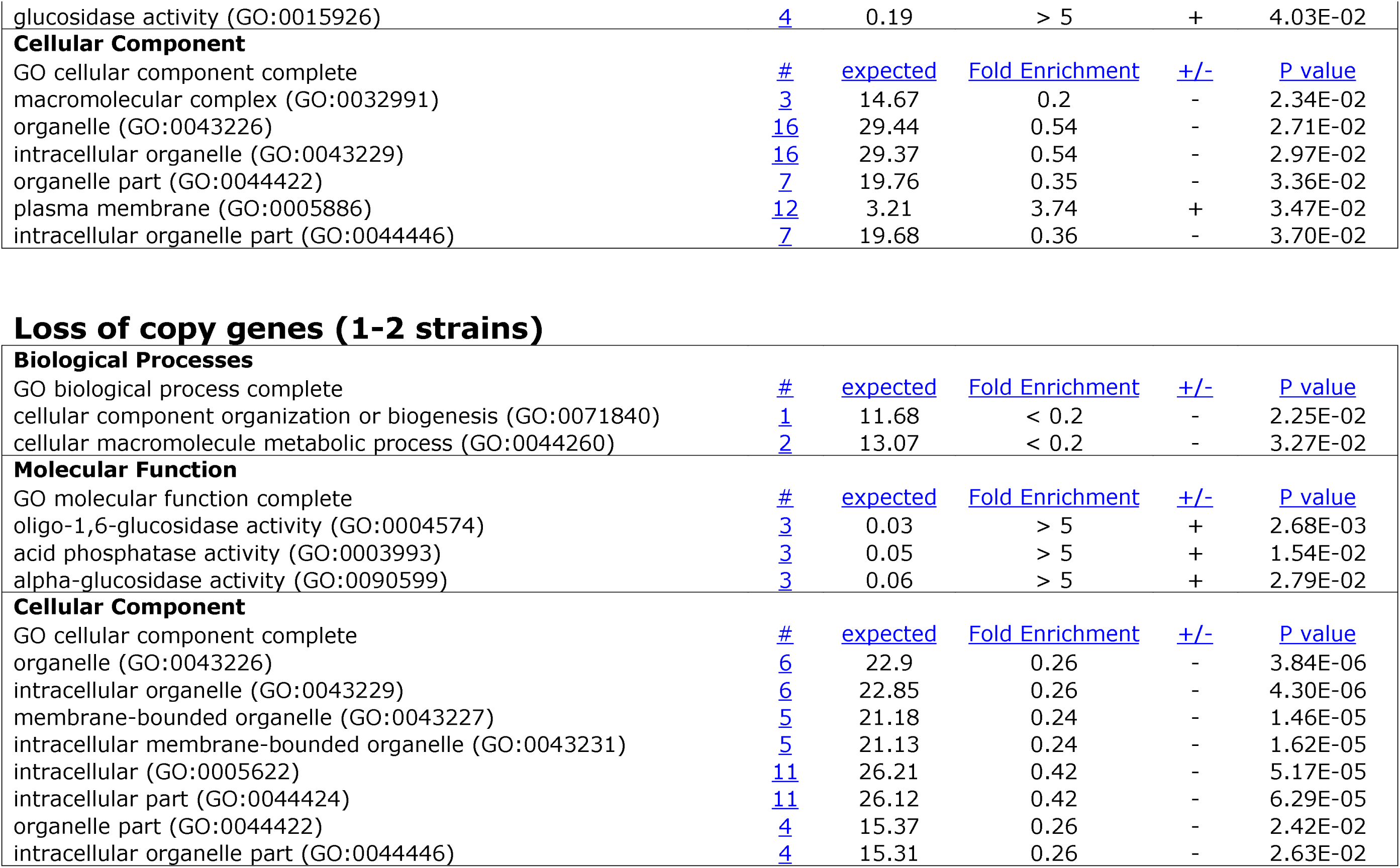
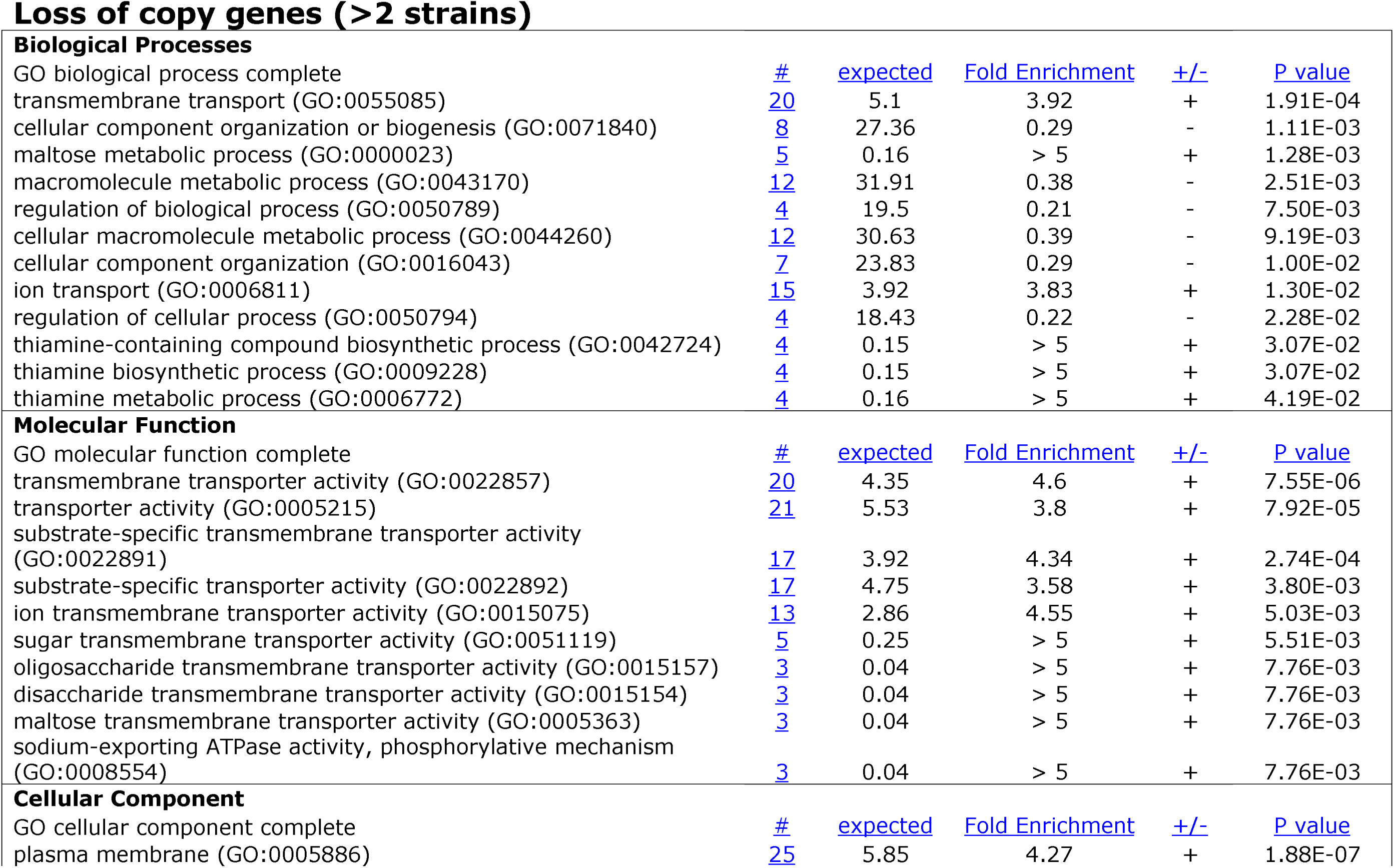
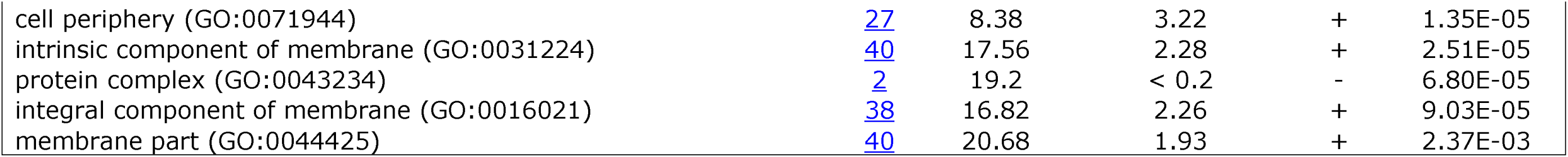
GO enrichment terms for genes that showed complete deletion, copy number gain, and copy number loss.

To observe patterns in gene copy number changes that are not deletions, we included only the 87 strains with no observed or suspected recent aneuploidies, where effects from those events are likely to dominate. 109 genes were significant for duplication [Table S4]. Rare duplications (63 genes found in 1-2 strains each) were just weakly enriched for thiosulfate sulfurtransferase activity, whereas common duplications (45 genes in 3-67 strains each) were enriched for genes in membrane transport [Table 1]. 119 genes were significant for copy number loss (excluding haploid strains) [Table S5]. Rare losses (35 genes; 1-2 strains each) were depleted for genes involved in organelle components, whereas common losses (82 genes; 3-68 strains each) was enriched for membrane components [Table 1]. Overall, 90/215 genes were members in 2 or more of the gene loss, gene copy number gain, and gene copy number loss lists, and likely represent genes that are highly variable in copy number among yeast strains.

## DISCUSSION

There has been a renewal in interest in natural populations of ‘wild’ *S. cerevisiae*, and the adaptive events that lead to commercial yeast strains (Barrio et al. 2006; Blondin et al. 2009; Muller and McCusker 2009b; Sicard and Legras 2011; Wang et al. 2012, Liti 2015; Peter and Schacherer 2015). A survey extending beyond the classic European strains used in lab or commercial environments showed that there is larger diversity in undomesticated *S. cerevisiae* strains from New Zealand (Gayevskiy and Goddard 2012). Another study on Chinese yeast strains found twice the amount of diversity as compared to previously surveyed strains, suggesting that Chinese yeasts may be a candidate source population (Wang et al., 2012). Geography thus plays a large part in wild yeast population structure, in stark contrast with commercial strains that have been spread across far reaching human communities through human activities (Goddard et al. 2010). To understand what differentiates virulent, commensal, wild, and commercially ‘safe’ strains in *S. cerevisiae*, it is necessary to understand the genetics of pathogenic yeast within this ever-growing context of non-clinical strains.

This task is hampered by the fact that a strains’ source substrate is a poor indication of whether it is truly a virulent, pathogenic clinical strain. The process of isolating *S. cerevisiae* colonies is often heavily biased towards strains that survive well in sugar rich conditions, and it is not always clear if *S. cerevisiae* is the dominant species occupying those substrates, or is even particularly well adapted to them (Goddard and Greig 2015). Earlier surveys of the *S. cerevisiae* population stucture and phenotypic diversity suggested that the shift from non-clinical to clinical yeast occurred multiple times, ruling out a single selective event that would have left clear genetic signatures. Instead, clinical strains isolated from human patients do not cluster into genetically distinct groups (Liti et al. 2009; Schacherer et al. 2009; Muller et al. 2011), and many remain genetically most similar to commercial strains (likely their source populations) rather than other clinical strains (de Llanos et al. 2006a,b; McKenzie 2008). The fact that multiple origins of clinical yeast strains have been found suggest a scenario where transitions are common, originate from multiple genetic backgrounds, and are unlikely to share a single adaptive strategy, if they are adapted to a pathogenic lifestyle at all. To fully study the transition of non-pathogenic yeast to a pathogenic lifestyle, either models are needed that can select for clinical behavior in non-clinical strains, to determine precise changes that might afford this change in lifestyle, or thousands of clinical and non-clinical yeast strains must be sequenced for a comprehensive analysis. Here, we start the process by re-sequencing 145 yeast strains, including 132 clinical isolates, from a variety of geographical locations and tissue substrates.

We find that earlier results hold over this larger set of clinical strains. Neighbor-joining tree and Structure results both confirm that many clinical isolates are more closely related to specific commercial strains instead of each other. We find evidence that there were likely independent pathogenic invasions from each of lab, sake, wine, and baking yeast strains. Within the rest of the clinical strains, there was no clear segregation of substrate tissue or geographical origin, and closely related strains often include isolates from entirely different tissue types and locations. It is possible that some of these strains may have been isolated from different tissues in the same patient, although we do not have access to the clinical data to confirm or refute that. It is also worth noting that due to considerations of comparability across datasets, we only used SNPs found and sufficiently covered in all datasets, which may bias the analysis towards relatively common polymorphisms.

Some clues about how yeast strains have adapted to drastic changes and stresses in new environments come from our understanding of how *S. cerevisiae* adapted to commercial fermentation (see Dequin and Casaregola 2011 for review). Functionally, whole genome or whole chromosome copy number changes can be beneficial under certain extreme conditions in yeast. Chromosome VIII, which carries the *CUP2* locus, consistently increases in copy number in a copper-rich environment (Whittaker et al 1988), and in general yeast appears to have the ability to maintain individual chromosomal copy numbers (Waghmare and Brusch 2005). Chromosome IX, the most often observed aneuploid in the clinical strains, is especially common in aneuploid state, even compared to other chromosomes (II, III and VIII) that are comparably stable as aneuploids (Zhu et al. 2012). These structural variants have been suggested to provide one-step short cuts when adapting to a harsh new habitat (Hittinger & Carroll 2007; Coyle and Kroll 2008; Rancati et al. 2008; Quero and Bond 2009; Stambuk et al. 2009, Muller and McCusker 2009b), and a similar process may be involved in clinical yeast adaptation going by the high frequency of polyploid and aneuploid strains. Association between the two also agrees well with the thinking that polyploid genomes are less stable than diploids/haploids, making them important drivers of rapid adaptation (Selmecki et al. 2015). More importantly, aneuploidies are highly unstable if not maintained by strong selective forces (Zhu et al. 2012; Kumaran et al. 2013; Zhu et al. 2014), and high aneuploidy rates could be indications that this is a selectable trait that may have been beneficial during the transition to pathogenic lifestyle, and would be worth detailed phenotypic studies, especially in clades such as the baker/commercial strains.

It must be noted that the majority of the clinical strains represented in our dataset were of North American and European origin. For a significant number of the clinical strains collected, we were unable to confidently identify the source population. Many of these strains are hybrids of known populations, but some may represent the genetic diversity of uncharacterized commercial or environmental *S.cer*. In addition, the lack of clinical strains that cluster with strains collected from Malaysia, Hawaii, and West Africa could indicate sampling bias. The role of increased human contact in opportunistic invasions may favor wine and baking strains, or it could indicate actual genetic predisposition of domesticated strains to transition to pathogenic lifestyles, perhaps precisely due to their genetic predisposition to easily adapt to drastically different carbon sources. More likely, it could be a combination of the two. Clinical strains thus present a selective view into a subset of medically relevant *S. cerevisiae*. More clinical strains collected from other parts of the world will help understand if clinical strains regularly colonize human hosts from non-commercial sources, or are mainly able to do so as a side effect of domestication prolonging exposure and opportunity.

It is undeniable that there is a large environmental shift when yeasts move into a pathogenic lifestyle. However, the corresponding genetic changes that resulted from these selective pressures are complex. While our knowledge of yeast diversity, complexity, and ecological traits are expanding, there remains a need for the whole genome deep sequencing of more yeast strains (Peter and Schacherer 2015), as well as the better documentation and classification of what exactly defines a strain as pathogenic and virulent. Better means of analyzing genomic data with variable ploidy across and within individuals will also be important in the study of *S. cerevisiae* isolates. While our dataset is limited in the types of analysis that can be conducted, in contrast to earlier studies on haploid sequencing data such as the 100-genomes from Strope et al. (2015), it provides information on a different scale and is complimentary to our understanding of the many levels of genetic variation available to *S.cer*. Such information, combined with human host traits and immune information, may provide the comprehensive picture required to tease apart yeast pathogenecity.

## MATERIALS AND METHODS

### Library Prep and Sequencing

Sample preparations were carried out in conjunction with strains prepared in Lang et al. 2013. Briefly, DNA was extracted using a modified glass bead lysis protocol. 500bp paired-end Illumina sequencing libraries were prepared at The Genome Institute, Washington University School of Medicine, and run on an Illumina HiSeq to an average depth of 80x coverage.

### Mapping and SNP Calling

The mapping pipeline was similar to that described in Zhu et al. 2014. Briefly, fastq files were mapped to the reference genome with bwa v0.5.9 (Li and Durbin 2009), sorted and indexed with samtools v0.1.18 (Li et al. 2009), and assigned strain ID with Picard tools v1.55. Duplicated read pairs were removed and remaining reads locally realigned with GATK v2.1-8 (McKenna et al. 2010). UnifiedGenotyper was used to call candidate variants across all samples simultaneously. The resulting VCF file was filtered for variants with coverage depth >8X, >2 reads and >15% of reads supporting alternative variant. Around 600kb of the genome – these regions were annotated in the SGD database as simple repeats, centromeric regions, telomeric regions, or LTRs (SGD project - http://downloads.yeastgenome.org/curation/chromosomal_feature/SGD_features.tab, downloaded 4th Aug 2012)– were excluded from analysis due to their susceptibility to mismapping and associated miscalls. For full list of commands, refer to Supplementary materials Zhu et al. 2014.

### Aneuploidy Calling, Gene Copy Number, and GO Term Analysis

Aneuploidy calling process was similar to method used in Zhu et al. 2014 and carried out by computing coverage depth of non-overlapping 1kb windows. Whole chromosome aneuploidies were called if average coverage of a chromosome differs more than 35% from other chromsomes in the same line (likelihood *P* < 0.001, χ^2^-test), and visually confirmed through coverage traces (see Supplemental Figures for examples). Gene copy number was estimated using normalized gene coverage against genome wide distribution, assuming binomial distribution of read depth. GO term analysis was conducted with g:Profiler (Reimand et al. 2011) and AmiGO 2 (Carbon et al. 2009).

### Neighbor-Joining Tree and Structure

Variants for the Liti strains (Liti et al. 2009) were identified according to an older version 2 of the yeast genome. Only sequenced SNPs from the Liti strains (not those inferred by algorithm) were converted to version 3 coordinates using LiftOver scripts from the UCSC genome browser website. SNPs called in both datasets with coverage in at least half of the strains within each dataset, were retained for NJ tree construction. The neighbor-joining tree was constructed using PhyML-3.1 (http://www.atgc-montpellier.fr/phyml), with missing data included as ‘N’. All tree figures were constructed with the online program Interactive Tree of Life (Letunic and Bork 2006). Structure (version 2.3.2) analysis was run with independent allele frequencies under an admixture model, for a 15,000 burn in period and 5,000 iterations of sampling.

---

Figure S1. Correlation between allele frequencies in non-clinical strains (X-axis) to allele frequencies in clinical strains (Y-axis).

Figure S2. Fst values (Y-axis) for alleles across the genome (X-axis).

Figure S3. Coverage plots for YJM264 across *S. cerevisiae* and *S. kudriavzevii* genomes. Chromosomes are lined in increasing order (X-axis), with total coverage in 1kb windows plotted (Y-axis).

Figure S4. Coverage plots for CBS2909 (a) and CBS2910 (b) across *S. cerevisiae* and *S. paradoxus* genomes. Chromosomes are lined in increasing order (X-axis), with total coverage in 1kb windows plotted (Y-axis).

Table S1. List of all strains included in this study.

Table S2. List of all fragments thought to be introgressed in strains CBS2909 and CBS2910, and genes that fall within these regions.

Tables S3-S5. List of all genes that showed complete deletion [Table S3], copy number gain [Table S4], and copy number loss [Table S5].

## Acknowledgements and Funding

Y.O.Z was supported by the A*STAR National Science Scholarship PhD. G.S was supported by R01 HG003328. D.A.P was supported by the NIH grants RO1GM100366 and RO1GM097415. We thank Greg. Lang, Patrick Gibney, and the WashU Genome Sequencing Center for generation of the DNA sequencing data. We thank John McCusker for sharing background knowledge on the clinical strains sequenced in this study.

